# Binding Affinity Ranking at the Molecular Initiating Event (BARMIE): An open-source computational pipeline for the rapid screening of chemical interactions with steroid receptors from many species

**DOI:** 10.1101/2025.02.05.636649

**Authors:** Fernando Calahorro, Parsa Fouladi, Alessandro Pandini, Matloob Khushi, Yogendra Gaihre, Nic Bury

## Abstract

A challenge in ecological risk assessment is identifying the chemicals that pose the greatest threat and determining which species are most vulnerable to them. To help address this, this study has developed an in-silico open-source tool called BARMIE (Binding Affinity Ranking at the Molecular Initiating Event) to rapidly predict the chemical binding affinity of steroid receptor proteins to synthetic steroids to identify potentially vulnerable species and chemicals of concern. BARMIE was used to screen 163 teleost fish glucocorticoid receptors (GRs) for binding to the natural ligand cortisol and to 10 synthetic glucocorticoid drugs (GCs) designed to interact within the ligand-binding pocket (LBP) of GRs. BARMIE identified species from the superorder Protacanthopterygii with high-affinity GRs to synthetic GCs (e.g. vulnerable species). . BARMIE was also used to screen binding profiles of compounds in the Medicine for Malaria Venture Global Health Priority Box to rainbow trout GRs (rtGR1 and rtGR2). Of the 178 compounds, 24 and 36 bind within the LBP of rtGR1 and rtGR2, respectively. For 30 of these compounds, transactivation activity was assessed at 1µM in the presence or absence of 1µM cortisol and confirmed 2 compounds with agonistic properties (e.g. chemicals of concern) that would require further in vitro and/or in vivo studies to assess the environmental risk. BARMIE can rapidly generate predicted binding affinities for 100’s of species and chemicals as a first screen in environmental risk assessment to provide information on which substances to prioritise in downstream tests.

## Introduction

Approximately 350,000 synthetic chemicals are produced globally, yet most have not undergone any human (HRA) or ecological risk assessment (ERA) [1]. Consequently, the impact of legacy and novel substances on human and wildlife health is likely underestimated. The evaluation of a chemical’s ecological risk requires information on mortality, growth, and reproduction, derived from toxicity tests, as well as on the chemical’s environmental fate, persistence and bioaccumulation properties. These toxicity tests are costly and time-consuming, and when fish are used, there are ethical concerns. Given the vast number of chemicals, it is impossible to rapidly generate sufficient toxicity data using current methods. In recent years, there has been a move to identify novel ways to provide the necessary toxicological information for ecological hazard and risk assessments that use rapid, high throughput in silico and in vitro methods, termed New Approach Methodologies (NAMs) [2–6].

All vertebrates have a similar steroid hormone/receptor-based endocrine system that regulates a plethora of physiological and developmental processes. These processes are primarily controlled by the binding of the natural steroid ligand to its receptor to initiate ligand-induced transactivation, transrepression, or stimulate other cellular signaling pathways [7]. These steroid receptor proteins (glucocorticoid, mineralocorticoid, androgen, estrogen, and progesterone) are highly conserved within the vertebrate subphylum [8].

The importance of the endocrine system is reflected in concerns about industrial chemicals, including pharmaceuticals, that are endocrine-active substances that perturb the action of hormones. In response to this concern, national and international governments implemented testing programs over 15 years ago to elucidate the endocrine-disrupting potential of synthetic and natural compounds [9,10]. These initially focused on the EATS (estrogen, androgen, thyroid, and steroidogenesis) modalities, but in more recent years, this has expanded to include non-EATS modalities [11] (e.g., glucocorticoid and progesterone receptors and non-steroidal receptors). The majority of NAMs’ research in the field of EDC has focused on HRA, with numerous in vitro and in silico tools available for the EATS modalities [12]. HRA aims to protect the individual; in contrast, ERA is far more ambitious and complex, seeking to protect populations of numerous species and maintain ecosystem function.

A good example of in silico based NAMS for ERA are platforms such as EcoDrug [8] and Sequence Alignment to Predict Across Species Susceptibility (SeqAPASS) [12], which use protein conservation across phyla to identify non-target species susceptible to drugs based on the presence of human or veterinary drug targets homologs. When these approaches are combined with other toxicological information, they yield a more comprehensive picture of the potential environmental impacts of chemicals. For example, SeqAPASS in combination with Genes to Pathway – Species Conservation Analysis (G2P-SCAN) uses network analysis, based on Adverse Outcome Pathway (AOP) information, to identify conserved Reactome [13] pathways and points of departure in toxic outcomes [14], and RASRTox (Rapidly Acquire, Score, and Rank Toxicology data) links the sequence information with toxicological databases (ECOTOX, ToxCast21) and QSAR models to develop tool for ranking chemicals for hazard assessment [15]. It is known that there are significant differences in the concentrations of chemicals that induce a toxic response between species within a taxa [16,17], thus a challenge is to increase confidence in predictive tools to identify sensitive species as well as chemicals of concern. One approach for endocrine active chemicals is to investigate the protein structural reasons for species differences in the interaction between chemicals and their receptors [18]. The hypothesis is that species with receptors with high binding affinity are likely to be affected at much lower concentrations of a chemical.

This study aimed to develop an in-silico tool for rapid screening of chemical binding to steroid receptors. The approach uses in-silico drug-discovery methodology [26] applied to predictive environmental toxicology. To demonstrate the utility of the tool we conducted two demonstration the first identifies vulnerable species based on the predicted receptor binding affinities to steroid drugs and the second to identify chemicals of concern within a library of non-steroidal chemicals. The study focuses on teleost fish glucocorticoid receptors (GRs) and the reasons for this are two-fold. Firstly, synthetic glucocorticoids, which are commonly used to treat various health conditions, are a group of drugs of growing environmental concerns, with 17% of the GCs being at concentrations in the aquatic environment predicted to pose a risk to fish [19]. Secondly, species differences in the binding affinity (Kd) and transactivation (EC50) activity for both natural and synthetic glucocorticoids (GCs) have been reported [20–25].

## Methods

### Computational model construction

The novel computational pipeline, Binding Affinity Ranking at the Molecular Initiating Event (BARMIE), uses open-source database APIs (UniProt [27], Chembl [28]) and software (OpenBabel [29], PyMol [30], and AutoDock Vina [31]). The code to estimate receptor binding affinities, a summary of the procedural steps, and user instructions are available at https://github.com/ParsaFouladi/Barmie, and a training video is provided at www.burylabs.co.uk. The pipeline was run at the University of Southampton HPC Iridis 6, and the example provided is specific to teleost fish GRs.

The pipeline can be adapted for use with other HPC architectures and other species and proteins.

### BARMIE overview

BARMIE uses fish protein receptor structures derived from AlphaFold [32] available in UniProt. These structures, in turn, are derived from the annotated fish genomes within the Ensembl database. Ensembl contains receptor isoforms (e.g., splice variants), and because very few isoforms have been characterized in fish [33], we did not remove these from our final analysis. When individual receptor structures are imported into AutoDock Vina, their orientations differ, and the box coordinates encompassing the ligand-binding pocket (LBP) for the docking experiment must be defined for each structure. To enable docking assessment for a large number of receptors and chemicals, the structures are automatically aligned using PyMol scripting so that the LBP location is in the same orientation for all proteins. The only manual step in the pipeline is defining the box coordinates for docking exploration of the LBP. This is set to a single reference protein, in this example, *Oncorhynchus mykiss* GR1 (accession # P49843), and is used within the BARMIE script for analysis of all receptors due to the three-dimensional alignment in PyMol. For the GRs utilized in this study, the docking box size was set to 20, 20, 20 Å, and the coordinates encompassed the LBP centred at X=5, Y=2, Z=-15.

Predicted docking binding affinity were estimated for 163 teleost GR proteins (SI 1 contains the Uniprot ID codes) in complex with the natural ligand cortisol (CHEMBL389621), the synthetic glucocorticoids beclomethasone (CHEMBL1586), clobetasol (CHEMBL1201362), dexamethasone (CHEMBL 384467), flumetasone (CHEMBL1201392), halcinonide (CHEMBL1200845), mapracorat (CHEMBL2103876), mometasone (CHEMBL1201404), prednicarbate (CHEMBL1200386), prednisolone (CHEMBL131), and triamcinolone (CHEMBL1451).

### BARMIE validation

AutoDock Vina uses a stochastic algorithm to explore ligand binding poses; thus, docking searches were run 5 times, and the average binding affinity is reported (SI 2 for cortisol). Docking simulations were run with exhaustiveness levels of 8, 32, and 128 to assess consistency of results (Table 1). We observed no difference among the exhaustiveness levels and report results from exhaustiveness 32 (Table 1).

**Table 1.**
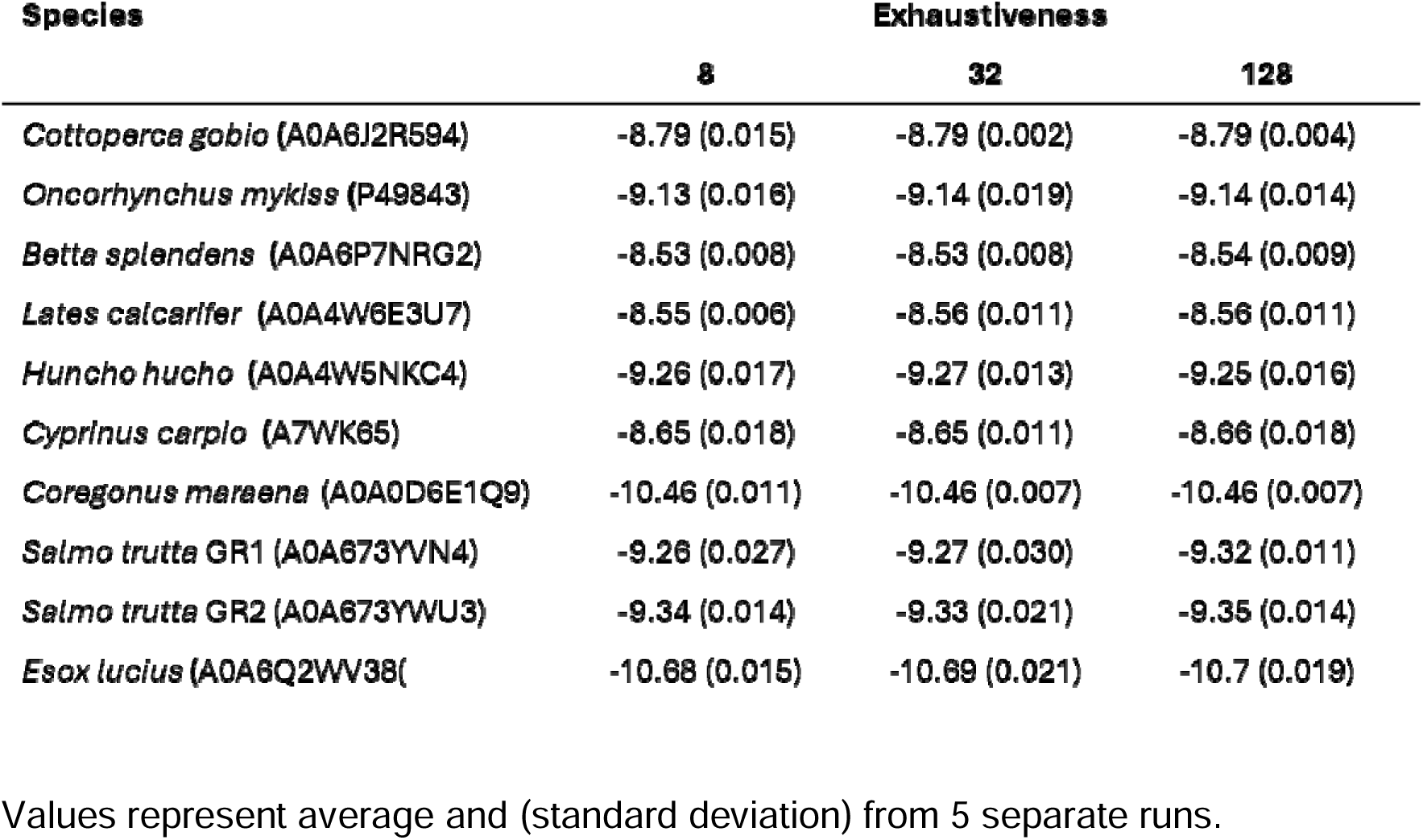
Predicted binding affinity generated in BARMIE at 3 Exhaustiveness values for 10 fish glucocorticoid receptors and cortisol. Values represent average and (standard deviation) from 5 separate runs.

| Species | Exhaustiveness |  |  |
| --- | --- | --- | --- |
|  | 8 | 32 | 128 |
| *Cottoperca gobio* (A0A6J2R594) | -8.79 (0.015) | -8.79 (0.002) | -8.79 (0.004) |
| *Oncorhynchus mykiss* (P49843) | -9.13 (0.016) | -9.14 (0.019) | -9.14 (0.014) |
| *Betta splendens* (A0A6P7NRG2) | -8.53 (0.008) | -8.53 (0.008) | -8.54 (0.009) |
| *Lates calcarifer* (A0A4W6E3U7) | -8.55 (0.006) | -8.56 (0.011) | -8.56 (0.011) |
| *Huncho hucho* (A0A4W5NKC4) | -9.26 (0.017) | -9.27 (0.013) | -9.25 (0.016) |
| *Cyprinus carpio* (A7WK65) | -8.65 (0.018) | -8.65 (0.011) | -8.66 (0.018) |
| *Coregonus maraena* (A0A0D6E1Q9) | -10.46 (0.011) | -10.46 (0.007) | -10.46 (0.007) |
| *Salmo trutta* GR1 (A0A673YVN4) | -9.26 (0.027) | -9.27 (0.030) | -9.32 (0.011) |
| *Salmo trutta* GR2 (A0A673YWU3) | -9.34 (0.014) | -9.33 (0.021) | -9.35 (0.014) |
| *Esox lucius* (A0A6Q2WV38) | -10.68 (0.015) | -10.69 (0.021) | -10.7 (0.019) |

For the fish GRs we used data from previous binding and transactivation activity experiments [34–37] to compare BARMIE binding affinity predictions to empirical dexamethasone binding data (Kd) and dexamethasone and cortisol log EC50 values for rainbow trout GR1 and 2 and their chimera clones and mutants [34,35], butterfly fish *Pantadon buchholzi* [36], sturgeon *Acipenser ruthensus* [36] and Common carp *Cyprinus carpio* [37]. In this experiment predicted GR protein structures were generated using AlphaFold v2.0 from the full-length GR amino acid sequence. These structures were fed into BARMIE to estimate predicted binding affinities. For the comparison of empirical binding data, the derived *K*d values were converted to ΔG using the Gibbs free energy relationship:

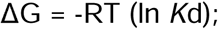

where R is the gas constant (0.0019872 kcal K^-1^ mol^-1^), T is temperature in Kelvin at room temperature (298K) and *K*d the empirical binding affinity in moles.

A comparison between the predicted BARMIE binding affinities and experimental values was done to confirm that the computational pipeline can produce comparable results. Linear regression models were calculated in GraphPad Prism v11.0.

### Screening of the Medicines for Malaria Venture Global Health Priority Box compounds

The Medicines for Malaria Venture (MMV) Global Health Priority Box (GHPB) chemical library was kindly supplied by Bristol-Myers Squibb Company and IVCC. As an example of how BARMIE could be used in a chemical screening programme we predict the binding affinity of the 178 compounds in the GHPB to rainbow trout GR1 (rtGR1) and GR2 (rtGR2) (SI 4). Thirty compounds (#1 to #30), with a binding affinity ≥ -7.5 kcal/mol and 10 compounds predicted to either bind with low affinity or outsideof the LBP (#31 to #40), as confirmed by visual inspection in PyMol. For this study a value of -7.5 kcal/mol was chosen because it is the average predicted binding affinity for mapracorat, a non-steroidal Selective Glucocorticoid Receptor Agonist (SEGRA) (SI 1). An in vitro transactivation assay [25] was used to assess agonist and antagonistic activity. Briefly, COS-7 cells were seeded into 96-well plates at a density of 10,000 cells per well and grown o/n in DMEM (Gibco) media containing 5% v/v FBS (Gibco) at 5% CO2 atmosphere and 37 °C. The following day, the cells in each well were transfected with 44.5 ng of the reporter plasmid PF31-LUC and 0.5 ng of the NanoLuc® control vector, PNL1.1. TK[Nluc/TK] (Promega) and five ng of the expression vector pcDNA 3.1(+) containing rainbow trout GR1 (rtGR1, UniProt # P49483) and GR2 (rtGR2, UniProt Q6RKQ3) supplied by Genscript Biotech using Escort™ IV Transfection reagent (Merck). The Global Health Priority Box compounds were resuspended in DMSO at a concentration of 10mM. After 24 hr, duplicate wells were incubated with either 1µM of each of the GHPB compounds or 1µM of GHPB compound and 1µM of cortisol in DMEM/F12 containing 5% charcoal-stripped FBS (Gibco) with a final concentration of 0.01% (v/v) DMSO. The cells were incubated for a further 24h. Then, firefly and Nluc luciferase activity was measured using Promega Glomax Navigator, following the methods described in the Dual-Glo protocol (Promega cat∼ N1630). Firefly luciferase activity was normalized to the Nluc activity and expressed as fold induction of the DMSO controls. For each experiment exposures were conducted in duplicate wells with experiments repeated three times except for compounds #36 to #40, which were performed in duplicate. Difference between the control values and each compound was assessed via a One-way ANOVA followed by a Dunnett’s post-hoc test using values from each well with significance attributed if p < 0.05. Statistical analysis was performed in GraphPad Prism v11.0

## Results and Discussion

A significant relationship was observed between the predicted binding affinities from BARMIE for rainbow trout GR1 and 2, their chimeras and mutants [34,35], *P. buchholzi* GR1 and 2a and *A. ruthenus* GR [36] with dexamethasone binding (Fig 1A; R^2^ = 0.52, p=0.0191) as well as EC50 values for dexamethasone and cortisol (Fig. 1B; R^2^ = 0.72, p<0.001, Fig 1b), with the two latter relationships included values for *C. carpi*o [37]. A previous study also demonstrated a good relationship between the predicted protein stability [ΔΔG (kcal/mol)] of a reconstructed ancestral CR and engineered mutations of this receptor and measured 11-deoxycorticosterone EC50 values (R^2^ = 0.78, [38]), as well as empirical dexamethasone binding and EC50 values [35]. The relationship between BARMIE’s predicted binding affinity and those derived manually was also significant (Fig 1C; R^2^ = 0.53, p<0.001). The results support the use of BARMIE as a rapid in-silico screening tool to predict chemical binding affinity to GRs. However, it is important to note that empirical data for both ligand binding and transactivation activity for fish GRs are sparse, and even sparser for non-GC chemicals, thus further studies are required to identify the extent of BARMIE’s applicability domain.

**Figure 1.**
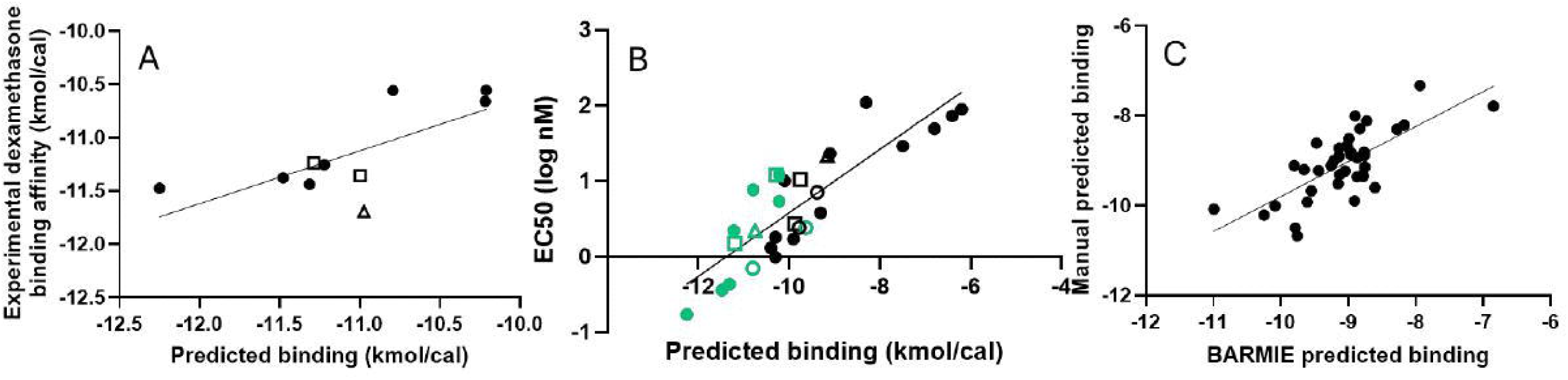
Comparison of predicted binding affinities and published (A) dexamethasone binding affinities (Kd); (B) log EC50 values for cortisol (black symbols) and dexamethasone green symbols and (C) a comparison of the predicted binding affinities generated in BARMIE and those derived manually for 40 GRs. Lines represent the linear regression analysis (A) R^2^ = 0.52, p=0.019; (B)R^2^ = 0.72, p<0.0001 and (C) R^2^= 0.53, p<0.001. The empirical data is from rainbow trout GR1 and 2, chimeras and mutations [solid circles] [34, 35], A*cipenser ruthenus* (open triangles) [36], *Pantodon buchholzi* GR1 and GR2a (open squares) [36] and *Cyprinus carpio* (open circles) [37] .

BARMIE predicted the binding affinity of 163 fish GRs to the natural ligand cortisol as well as synthetic GCs that are designed to interact within the LBP of GRs (Fig. 2). From the in silico binding affinity screen, halcinonide (Fig. 1) may be considered the most potent GC to interact with the fish GRs, whereas prednicarbate (Fig. 1) is the least potent. For all GCs, the screen shows that species in the order Protacanthopterygii (*Esox lucius, Salmo trutta, Coregonus mareana*) possess GRs with high binding affinity for the synthetic GC (Fig. 1A). In addition, 55% of the five most sensitive GRs belong to the Protacanthopterygii, with the species Northern pike (*Esox lucius*) ranked 1st for 7 of the 11 GCs tested (Table SI 1). This exercise illustrates how sensitive species may be identified with the data available, but there is a caveat. There are only 86 fish genomes have been annotated out of the potentially 30,000 teleost species [38], and of these genomes, a high proportion, 8%, are Protacanthopterygii, thus they are over represented in our analysis and other species maybe more sensitive. Looking forward, the number of species genomes sequenced is rapidly expanding with the Earth BioGenome Project Network [39] setting an ambitious target of “… characterizing the genomes of all of Earth’s eukaryotic biodiversity over a period of ten years”. This has the potential to provide a wealth of genetic information for in silico approaches in ecotoxicology.

**Figure 2.**
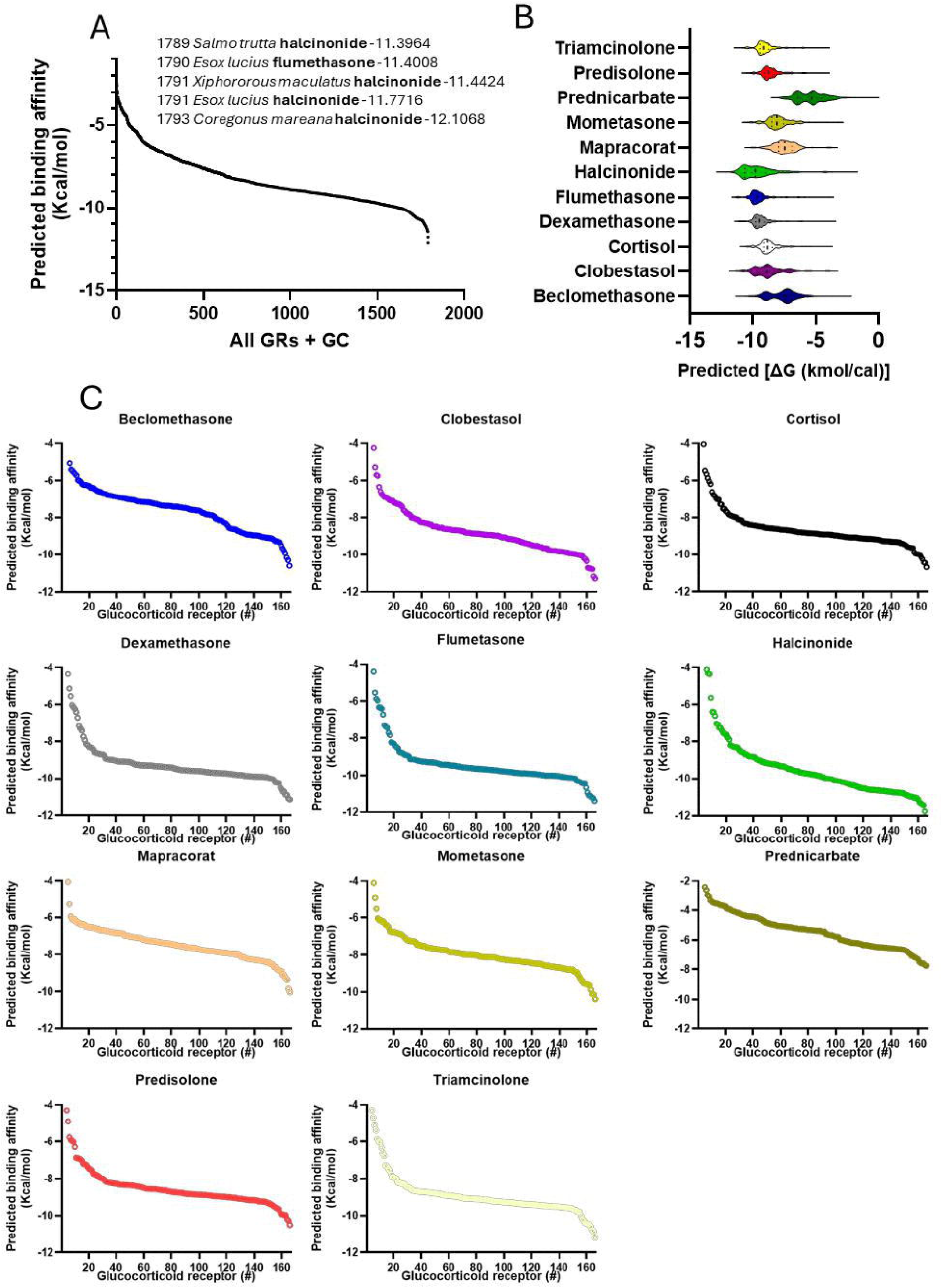
A. The predicted binding affinity for all chemical and species combinations, the inset list provides the top 5 species + chemical binding affinities. B Violin plot showing spread of binding affinities per GC. C. Binding affinities for the 163 teleost fish glucocorticoid receptors per glucocorticoid (see supplementary information for further species values).

BARMIE was initially used to provide docking information for steroids designed with a backbone comprising 17 carbon atoms arranged in 4 rings to interact with the LBP of GRs. Industrial chemicals may not have this structure, but they may nonetheless dock into the LBP to act as receptor agonists or antagonists. The binding of a compound into the LBP is a necessary first step in activating the protein, but binding of non-steroidal chemicals may not lead to a biological response . For this to occur communication between the LBP and the transactivation function site, termed AF2 in helix 12 of GRs, is necessary to enable the recruitment of the transcriptional machinery for receptor activation and gene expression [40]. The signalling pathway between the LBP and the AF2 site begins with hydrogen bond formation between the ligand and key amino acids within LBP [41], which induces the necessary protein conformational changes for activation. Despite the acknowledged complexity of the structural changes essential for receptor function, in-silico tools are often used in the initial stages of a drug discovery pipeline to identify promising leads to reduce the number of compounds to be tested in downstream transactivation assays [26].

Taking the traditional drug discovery approach, we suggest a workflow using BARMIE as the first stage in identifying potential leads for chemicals of concern (Fig 3). To screen the 178 GHPB compounds we focused on the rainbow trout GRs because these were used in the validatory exercise. We identified 24 and 36 of these compounds that bind within the LBP of rtGR1 and GR2 with a predicted binding affinity of ≥ -7.5 kcal/mol. The highest binding affinity observed for these compounds was -8.908 and -9.953 kcal/mol for GR1 and GR2, respectively, not too dissimilar to BARMIE predicted cortisol binding affinities to rtGR1 and GR2 of -9.383 kcal/mol and -9.98 kcal/mol, respectively. From these, 30 lead compounds were selected to assess agonist and antagonist behaviour at a concentration of 1µM in the presence or absence of 1µM cortisol, as well as 10 compounds that were predicted either not to bind or weakly bind to the receptors. Only a handful of compounds showed significant agonist activity (Fig. 4), and activity was substantially lower than the 100-fold increase observed with cortisol (Fig. 4). The reason for this low activity may be that the one µM concentration is insufficient to elicit a response in many of these compounds, as they are not designed to mimic the structure of endogenous cortisol. The concentration was chosen because it reflects elevated plasma cortisol levels in fish [42] and also corresponds to the maximum rtGR transactivation activity for cortisol [20,25].

**Figure 3.**
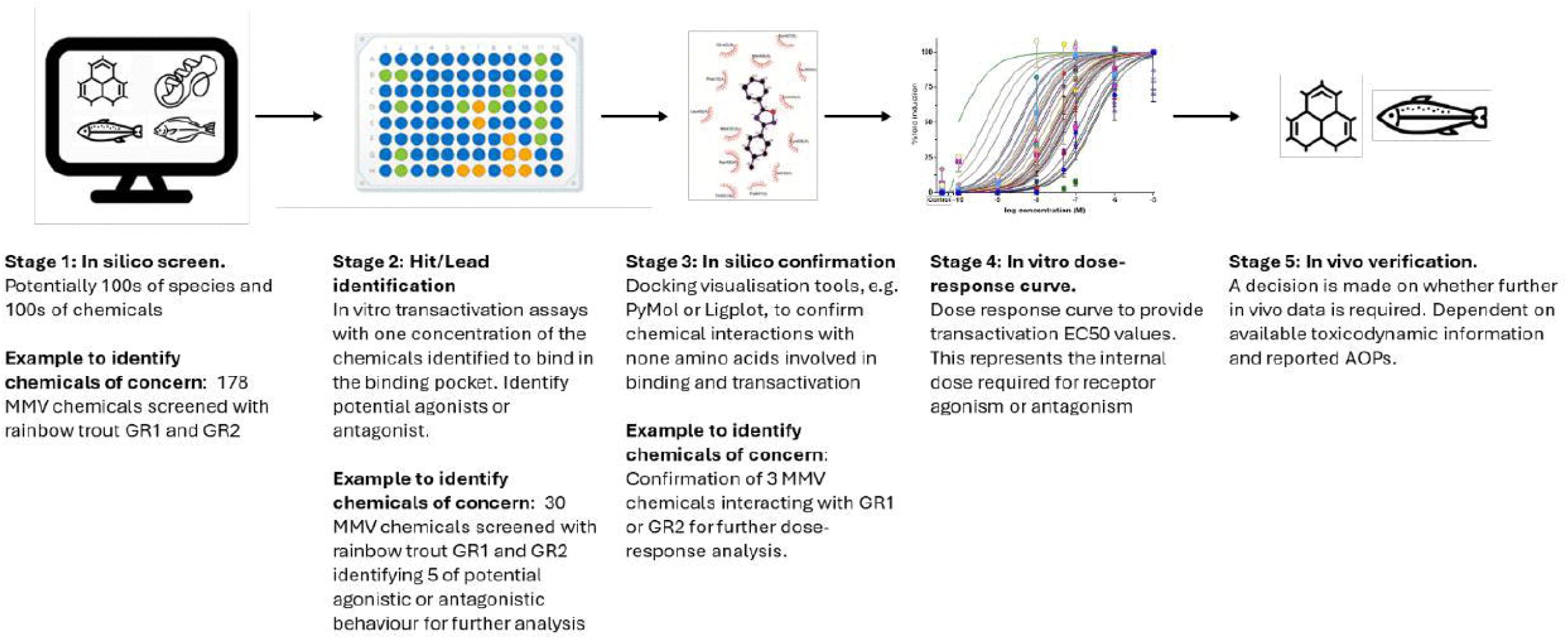
A suggested workflow for use of BARMIE to identify potential chemicals of concern.

**Figure 4.**
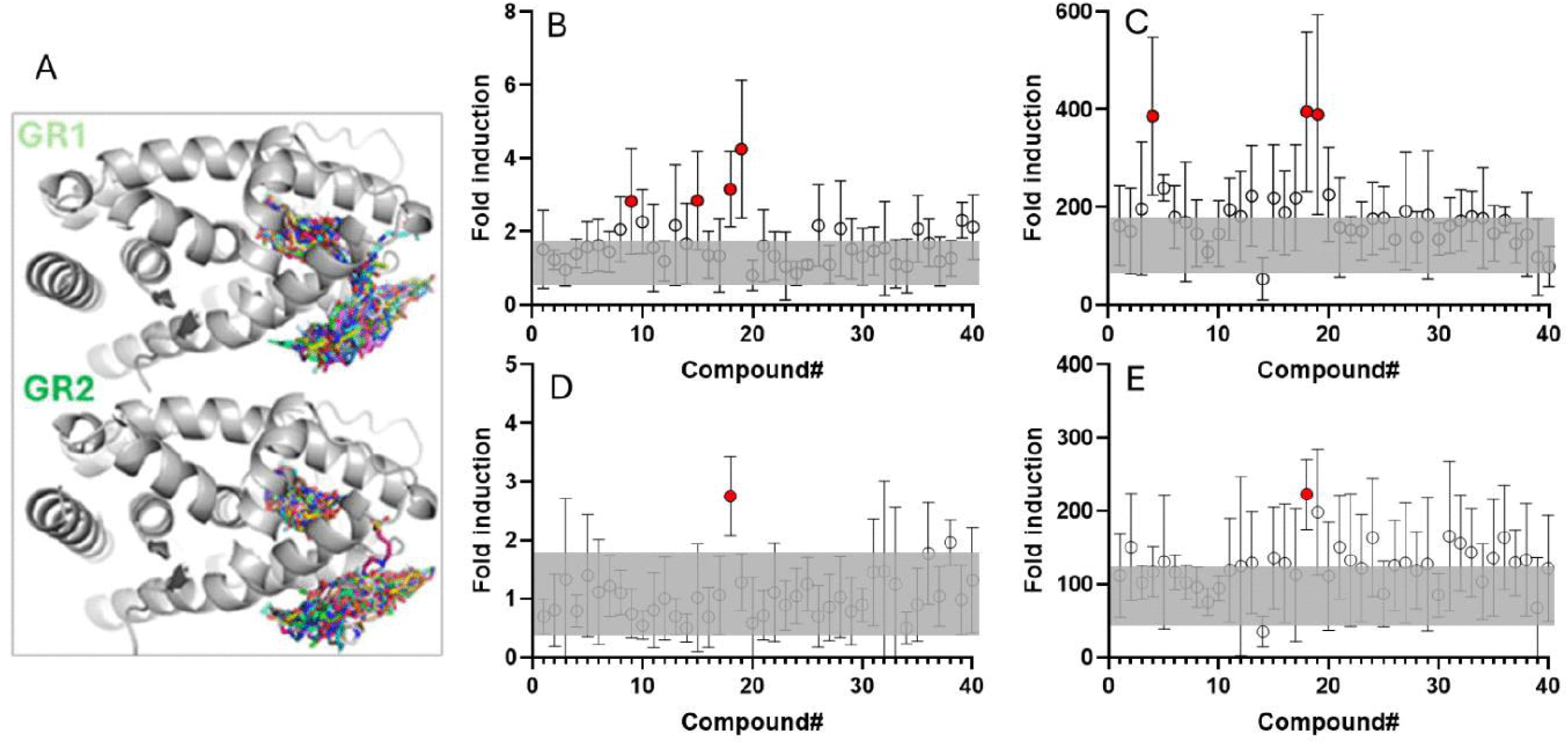
A. a composite image of the ligand binding profiles of the MMV Global Health Priority Box from BARMIE. B – E. Fold induction for rtGR1 (B & C) and rtGR2 (D & E) relative to DMSO controls for compounds #1 to #30 predicted to bind with the ligand bind pocket (LBP) with a binding affinity of ≥-7.5 kcal/mol and #30 to #40 for those compounds predicted not to bind with the LBP in absence (B & D) or presence of 1µM cortisol (C & E). The grey rectangles represent the range (average ± SD) of values from the DMSO controls (B & C) and 1µM cortisol alone (D & E). The red circle indicates those compounds where values are significantly different from the controls (One-way ANOVA with Dunnett’s post-hoc -test, P<0.05). Values represent average of 3 separate experiments, with duplicate wells per experiment, ± SD, except #36 to #40 where the values represent repeat experiments. See SI 4 for list of compounds and predicted binding affinities.

Two compounds from the screen are of potential concern. #18 (3-(4-methylphenyl)-5-phenyl-1,2,4-oxadiazole), showed significant agonist activity for both rtGR1 and rtGR2 in the absence and presence of cortisol and #19 (3-(3-methyl-2-pyridinyl)-5-phenyl-1,2,4-oxadiazole) significant activity in rtGR1 (Fig 4). Three others, # 4, 9,15 stimulated GR1 but only in one of the agonist screens (e.g. either in absence or presence of cortisol).Both #18 and 19 are 1,2,4-oxadiazole compounds known to have biological activities, including estrogen receptor antagonism [43]. None have been reported as GR agonists in fish, and we are not aware of any reports of their concentrations in the environment. Thus, these would be candidates for further studies in a risk assessment workflow as outlined in Fig 3.

Understanding the reasons for non-steroidal chemical agonism or antagonism is complex. For example, from the amino acid interaction plots, compounds #18 and #19 both form hydrogen bonds with Asn539 of rtGR1(Fig. 5), an essential amino acid in the docking of the natural ligand cortisol into the LBP [41] (Fig. 5), which is involved in the initiation of the signal to enable transcriptional machinery recruitment.

**Figure 5.**
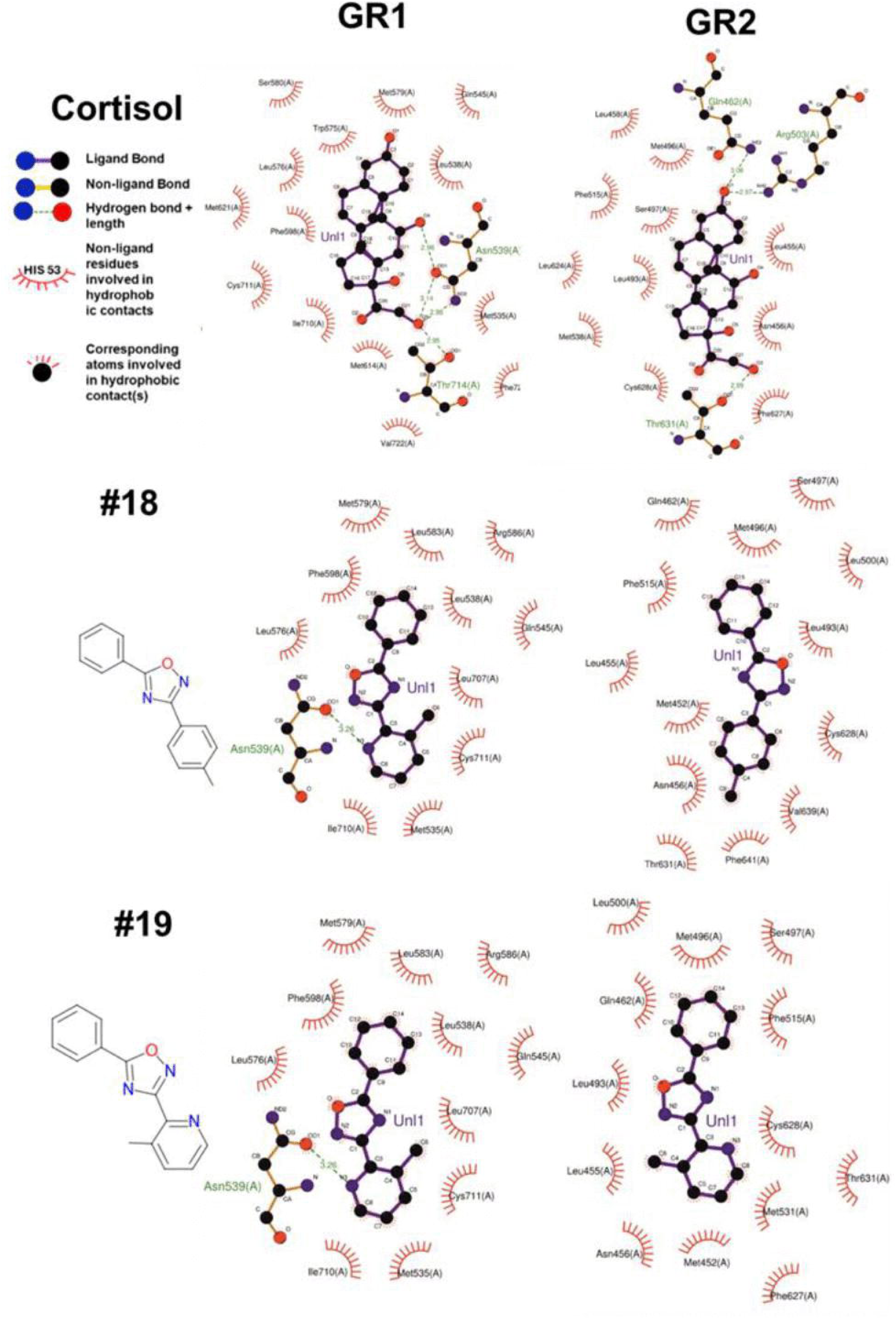
The amino acid interactions between compound #,18,19 and cortisol with rainbow trout GR1 and 2, as predicted by LigPlot [48].

However, #18 also stimulates rtGR2, and there is no predicted hydrogen bond formation with those amino acids known to interact with cortisol and potential potency is associated with Van der Waal interactions, as is the case with the GC fluticasone furoate in the human GR [45]. Further work is required to better understand the link between non-steroidal chemical interactions with fish steroid receptors and their agonist or antagonistic behaviour.

## Conclusions

BARMIE provides an open-access tool for rapid screening of synthetic drug ligands and industrial chemical binding affinities to proteins. The current study used glucocorticoid receptors as an example, but this can be expanded to other steroid and non-steroid receptors, as well as other proteins at the molecular initiating event (MIE) of an Adverse Outcome Pathway (AOP) [46]. Validation of BARMIE is promising, indicating a link between predicted binding and gene activation, but we must stress caution because for fish GRs there are only a handful of empirical datasets available for this validatory exercise. We suggest a way in which BARMIE can be incorporated within a chemical environmental risk assessment workflow (Fig 3) that is similar to the in-silico approaches used in drug discovery. In this scenario BARMIE would be used as a first screen to identify potential leads, these leads are then screened for agonists or antagonistic properties before more extensive (and costly) dose-responsive curves to determine EC or IC50 values, and or in vivo studies, are conducted.

## Supporting information

Supplemental Ivformation 2

## Acknowledgements

We would like to thank Medicines for Malaria Venture (MMV) Global Health Priority Box (GHPB) chemical library supplied by Bristol-Myers Squibb Company and IVCC for the chemical libraries.

## Author Contribution declaration

PF: software; methodology; review and editing.

FC: software; validation; methodology; data curation; formal analysis; review and editing.

YG: validation; formal analysis; review and editing. AP: software; supervision; review and editing.

MK: conceptualisation; software supervision, funding acquisition; review and editing.

NB: conceptualisation; funding acquisition; resources; supervision, project administration; first draft and revisions.

## Data availability statement

BARMIE code and installation instructions are available on Github https://github.com/ParsaFouladi/Barmie.

## Supporting information

Supplementary Information 1 – Excell spreadsheet with predicted binding affinity of various synthetic glucocorticoids and cortisol to fish glucocorticoid receptors.

Supplementary Information 2 – Five repeat runs with BARMIE for cortisol.

Supplementary Information 3 – Summary of predicted binding affinities to the MMV GHPB compounds to rainbow trout GR1 and GR2.

